# Arm locomotor behaviour affects motor unit discharge characteristics in the stationary leg

**DOI:** 10.64898/2026.08.17.745355

**Authors:** Benjamin M. Nazaroff, Emma R. Mitchell, Gregory E.P. Pearcey

## Abstract

Persistent inward currents (PICs), which are facilitated by monoaminergic inputs such as serotonin (5-HT), amplify synaptic drive and strongly influence motoneuron excitability. Although rhythmic locomotor activity increases serotonergic drive in animal models, its effects on intrinsic motoneuron properties in humans remain unclear. We examined whether rhythmic arm cycling alters motoneuron excitability of the non-exercising tibialis anterior during submaximal contractions. Twelve healthy adults (8 males, 4 females) performed triangular isometric dorsiflexion contractions at 25% and 50% MVC under four conditions: resting arm (CONTROL), finger tapping (TAP), arm cycling at 50–60 RPM (LOW), and arm cycling at 80–90 RPM (HIGH). Motor unit activity was identified from high-density surface electromyography that was decomposed into spike trains. Recruitment thresholds of identified and tracked motor units were consistent across conditions, but ΔF (i.e., an estimate of the PIC-related contributions to motor unit discharge) decreased during high-cadence arm cycling at stronger contraction intensities, which may reflect either reduced neuromodulation and/or increased or altered patterns of inhibition. In contrast, ascending discharge rate modulation deviated from linearity to a greater extent (i.e., brace height was larger) during both low- and high-cadence cycling, indicating greater neuromodulatory influence on the ascending discharge rate pattern. Self-sustained discharge was also elevated during cycling tasks, reflecting prolonged motor unit discharge. Taken together, these findings suggest that rhythmic activity of the arms modulates the discharge characteristics of motoneuron pools in the legs via unique combinations of excitatory, neuromodulatory and inhibitory inputs, which advances our understanding on the mechanisms of interlimb neural coupling.

## INTRODUCTION

All motor output occurs via activation of motor units, which are comprised of spinal motoneurons and their innervated muscle fibers (Heckman & Enoka, 2012). Motoneurons receive both ionotropic excitatory and inhibitory inputs, as well as monoaminergic inputs that are neuromodulatory in nature that dramatically influence their input-output properties (Binder et al., 2020; Heckman et al., 2009). Serotonin (5-HT) is one of the primary monoamines released from the brainstem that alters the discharge properties of motoneurons (Johnson et al., 2017) by facilitating persistent inward currents (PICs), which provide gain control via amplification and prolongation of synaptic inputs (Khurram et al., 2021). Monoamine (i.e., 5-HT) release is highly diffuse and is likely to play a role in mediating state changes of the excitability of motoneurons (MacDonell et al., 2015; Power et al., 2010). Locomotion, in particular, increases the discharge rate of descending serotonergic neurons that release 5-HT onto motoneurons throughout the spinal cord, which is proportional to the speed of walking in cats (Jacobs et al., 2002). The contribution of 5-HT to state changes in human motoneuron excitability, however, remains poorly understood.

Activity in one limb can influence excitability and motor output in another. For example, rhythmic upper limb movement has been shown to modulate the excitability of spinal reflex pathways in the legs, even when the legs remain stationary (Frigon et al., 2004; Loadman & Zehr, 2007; Pearcey & Zehr, 2019). This interlimb coupling is thought to arise from shared networks in the spinal cord, such as central pattern generators (CPGs) and interneuronal pathways (Zehr et al., 2016), and from changes in descending drive and sensory feedback during locomotor activity (Katz, 2016). Whilst most research on arm and leg neural interactions has focused on reflex modulation or corticospinal excitability, less is known about how rhythmic upper limb movement influences estimates of intrinsic motoneuron properties in the lower limb, such as those shaped by neuromodulators such as 5-HT. Understanding these effects could reveal how rhythmic movement in one part of the body sets the state of neural excitability of remote motoneuron pools, with potential applications for rehabilitation strategies that use arm cycling or other rhythmic tasks to prime the nervous system (Kaupp et al., 2018; Porter et al., 2025)

Despite research in reduced preparations showing that locomotor activity increases monoaminergic drive to motoneurons (Feraboli-Lohnherr et al., 1999; Jacobs et al., 2002; Veasey et al., 1995), and motor unit discharge rates being enhanced during arm cycling compared to an intensity-matched isometric contraction (Basile et al., 2025), rhythmic arm cycling effects on estimates of the contribution of PICs to voluntary contractions in the leg remain unknown. Remote, isometric upper limb contractions enhance estimates of the contribution of PICs to the discharge of leg motoneurons (Orssatto et al., 2022), but little is known about rhythmic arm movement effects, which engage distinct spinal and supraspinal pathways (Frigon, 2017). Recent work has shown that the VibStim protocol (i.e., a method utilising neuromuscular electrical stimulation and tendon vibration of the triceps surae to estimate PIC contributions to involuntary force output) is unaltered with simultaneous arm cycling (Alahmari et al., 2026). The authors suggested that the lack of an effect may be attributed to an insufficient arm cycling intensity to modulate motoneuron excitability or ceiling effects of the stimulation intensity. This is further confounded by the fact that rhythmic arm movement may generate inhibitory spinal commands that suppress lower limb reflexes, potentially counteracting additional depolarizing currents from PICs during arm cycling (Dragert & Zehr, 2009).

We further explored this concept and exploited the unique combination of excitatory and inhibitory influences onto motoneurons induced by rhythmic motor output of the arms compared to static tasks. More specifically, the purpose of this study was to determine whether rhythmic arm cycling alters the motor unit discharge characteristics in the stationary (i.e., non-exercising) tibialis anterior (TA) during low- and moderate-intensity isometric contractions. We hypothesized that locomotor activity in the arms would cause diffuse 5-HT release, thereby facilitating the contribution of PICs to the discharge of motor units in the stationary legs.

## METHODOLOGY

### Participants and Ethical Approval

Seventeen healthy adults were recruited for this study; however, five were excluded because their recorded data was not usable due to either poor pulse-to-noise ratios of identified motor unit spike trains or the inability to adequately match the target profile of dorsiflexion force during arm-cycling trials. After these exclusions, twelve participants (8 males; 25 ± 7 years, 4 females; 22 ± 2 years) were included in the analysis. Participants had no history of neuromuscular disease, recent muscular injury, or other abnormalities that would prevent them from completing tasks described below. The study’s experimental procedure was in accordance with the Helsinki Declaration, except for the fact the study was not registered, and all protocols were approved by the Interdisciplinary Committee on Ethics in Human Research at Memorial University of Newfoundland (ICEHR no. 20231547-HK).

### Experimental Set Up

Participants performed either rested their arms, tapped to a metronome or performed arm cycling on an Upper Body Ergometer (Monark Rehab Trainer 881 E, MONARK, SWEDEN) while performing dorsiflexion contractions. Cycling cadence was monitored by a digital display, and participants were asked to maintain a range of revolutions per minute (RPM). Cycling intensity, measured in watts (w), was adjusted by turning a knob that adjusts the tension on the flywheel of the cycling ergometer. The cycling wattage was maintained at 5 w and monitored on the built-in display on the ergometer. Participants maintained a constant cadence of 50-60 RPM, and 80-90 RPM for the LOW and HIGH conditions respectively, and the tensioner was adjusted at each cadence accordingly to meet the desired cycling wattage. The cycling ergometer was positioned at a distance and height from each participant such that were able to fully extend their elbow joint at the “3 o’clock” position, as depicted in Figure 1.

**Figure 1.**
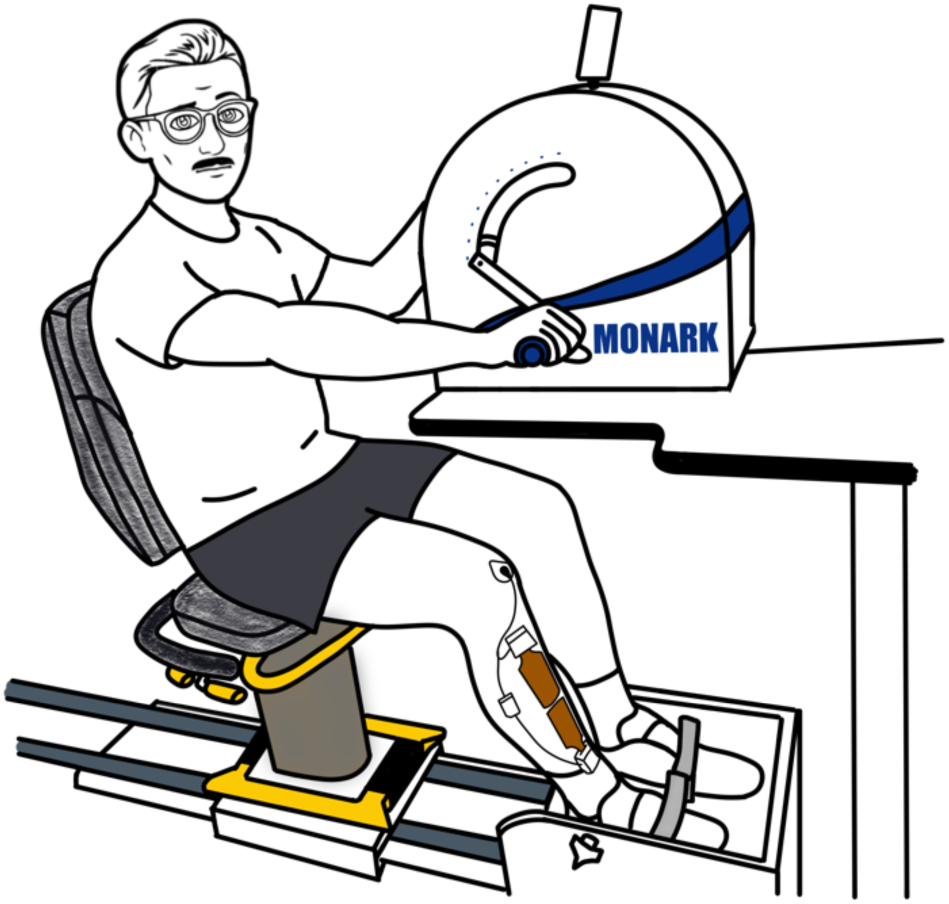
The experimental set up of participants during the arm cycling conditions.

### Experimental Protocol

Participants performed a minimum of two maximum voluntary isometric dorsiflexion contractions (MVCs) for 3 seconds, and then 90 seconds of rest was allotted between each MVC. A third MVC was performed if there was >5% difference between the first two MVCs. During MVCs, participants received verbal encouragement to ensure maximal muscle contraction, and maximum values were used to normalize subsequent submaximal trials.

Following MVCs, participants performed a series of submaximal ramp dorsiflexion contractions, while also completing rhythmic and non-rhythmic tasks in the upper limbs. The ramp contractions were 20 seconds in length; the first 10 seconds was a linear ramp to a specific percentage of their MVC and once the peak of the ramp contraction was reached, participants immediately started to perform a 10 seconds ramp down to rest. These ramping contractions were performed to a peak of 25 and 50% MVC. There were four conditions in which participants performed submaximal contractions: 1) ***CONTROL:*** submaximal dorsiflexion ramping contractions performed with no upper body movement; 2) ***TAP***: submaximal dorsiflexion ramping contractions performed while participants moved their index finger right to left when cued by a pseudorandom beep from a prerecorded audio file; 3) ***LOW***: submaximal dorsiflexion ramping contractions performed while arm cycling at 50-60 RPM on the Monark Arm Cycling Ergometer; and 4) ***HIGH***: submaximal dorsiflexion ramping contractions performed while arm cycling at 80-90 RPM on the Monark Arm Cycling Ergometer.

Participants began by performing two ramp contractions each at 25% and 50% MVC in the CONTROL condition. After a 5-minute washout period, participants performed two ramp contractions at 25% MVC and 50% MVC under the TAP condition. Following another 5-minute washout period, participants performed two 25% and 50% dorsiflexion ramp contractions under the LOW cycling condition, followed by another 5-minute washout period, after which they concluded the protocol by performing a final two 25% and 50% dorsiflexion ramp contractions under the HIGH cycling condition. In total, each participant performed 16 submaximal dorsiflexion contractions. An experimental timeline is outlined in Figure 2. A minimum of 30 seconds of rest was allotted between each contraction within a condition. This experimental design was intentionally not randomized because Orssatto et al. (2022) have reported that performing handgrip contractions prior to contracting the TA increases ΔF in the leg, whereas there are not changes during a control (i.e., 30-seconds of rest) condition. These findings therefore shaped our protocol: any trial involving upper body activity was performed after the CONTROL condition to avoid any potential carry-over effects. Furthermore, the three upper body conditions (TAP, LOW, and HIGH) were ordered by increasing intensity in the upper body, with lower-cadence trials conducted first. This ordering was based on the expectation that voluntary drive would progressively increase with higher-cadence arm cycling, and was designed to prevent high-cadence trials from influencing estimates of the contribution of PICs in subsequent lower-cadence conditions.

**Figure 2.**
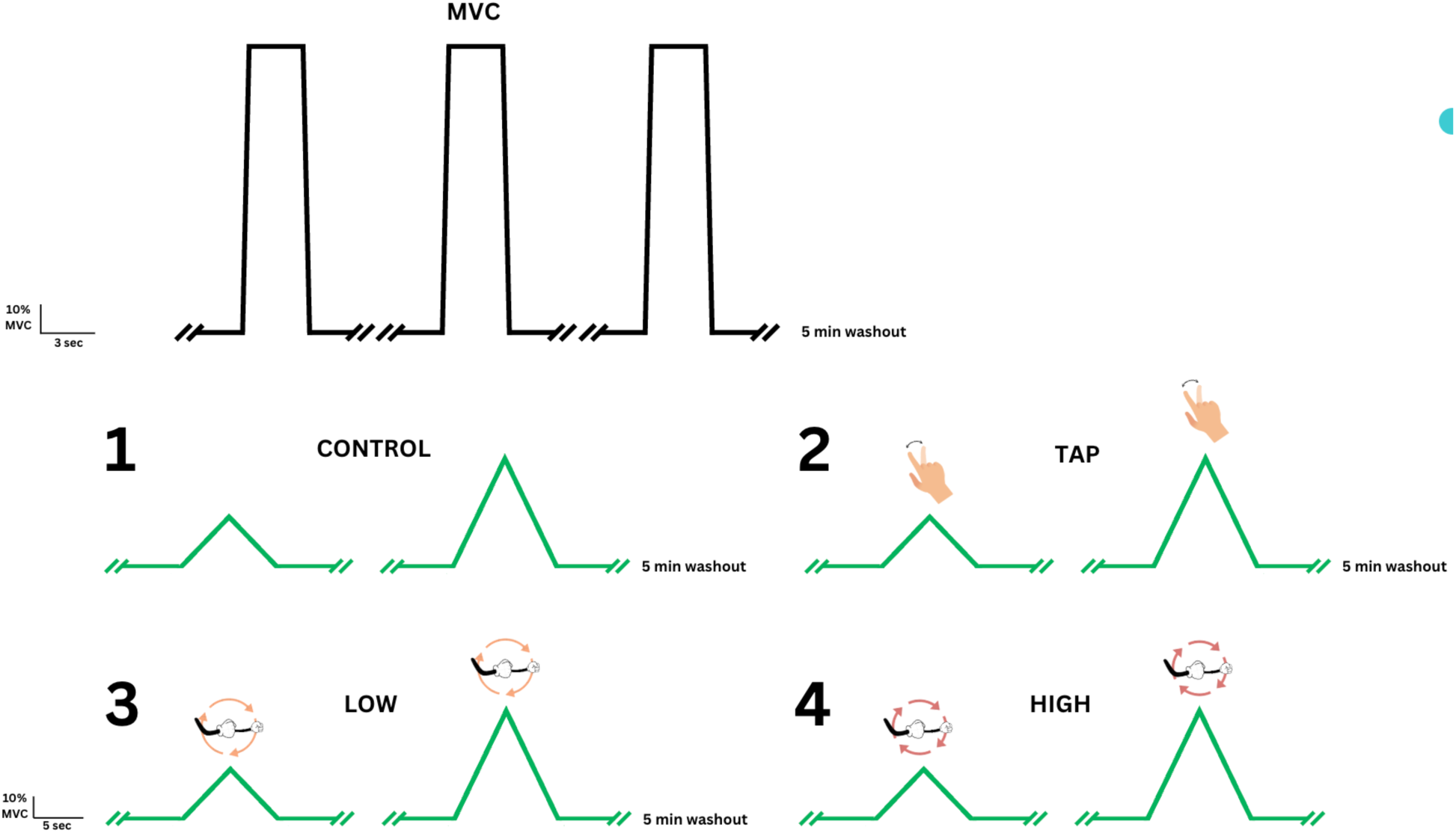
Experimental timeline of protocol. Conditions are labelled with numbers and the order in which they were performed.

### Force and High-density surface electromyography (HD-sEMG) Recording

Force and HD-sEMG signals were used to assess the volitional motor output of the tibialis anterior. Participants were seated in a chair with their feet secured to a custom footplate with built in strain gauges (Omegadyne model 101-500) that was used to measure ankle dorsiflexion forces with their right ankle at ∼95° while participants were seated in an adjustable chair such that their hip and knee joint were each in 100° of flexion. Force was sampled at 2048 Hz and smoothed offline with a 20 Hz low-pass filter (fifth-order Butterworth).

Prior to placing electrodes on the muscle of interest, the site was shaved and abraded with a conductive paste (Nuprep Skin Prep Gel, Weaver and Company, USA). Two 32 channel HD-sEMG arrays (4 x 8, 10mm I.E.D., ReC Bioengineering Laboratories and LISiN, Turin, Italy) were placed longitudinally over the tibialis anterior to sample motor units of the ankle dorsiflexors. Two Ag/AgCl ground electrodes were placed on the patella and the lateral malleolus of the right ankle for the proximal and distal electrodes, respectively. The HD-sEMG signals were collected in monopolar mode, sampled at 2048 Hz, amplified x150, and band-pass filtered at 10-500Hz using miniature wireless signal amplifiers (MEACS, ReC Bioengineering Laboratories and LISiN, Turin, Italy). A 1-second TTL pulse was transmitted to the wireless amplifiers to synchronize HD-sEMG and force signals.

### Data Analysis

#### Motor unit decomposition

After collection, monopolar EMG signals from each channel were bandpass filtered at 10-500 Hz utilizing a second-order Butterworth filter. Signals were then visually inspected, and channels with substantial artifacts or noise were removed. Individual motor unit spike trains were then decomposed with a convolutive blind-source separation algorithm (Holobar et al., 2014; Holobar & Farina, 2014) implemented in the DEMUSE software tool (v5.01; The University of Maribor, Slovenia). Extraction parameters were set to 100 iterations with a maximum coefficient of variation of 50%. This decomposition procedure has been extensively validated using experimental and simulated signals with respect to identifying motor unit discharge times over a range of contraction intensities (Holobar and Farina, 2014; Holobar et al., 2014).

After decomposition, motor unit spike trains were visually inspected and manually edited using the CKC inspector within the DEMUSE software tool to correct minor errors introduced by the decomposition algorithm. Well-validated local re-optimization methods were applied to improve the accuracy of motor unit spike trains, following techniques similar to those used in recent studies (Borzuola et al., 2023; Hug et al., 2021; Škarabot, Thomason, et al., 2025). The inverse of the interspike interval were computed to determine instantaneous discharge rates of each motor unit spike train, which were then smoothed using support vector regression (Beauchamp et al., 2022) with custom-written MATLAB scripts. Initial, peak, and final discharge rates were extracted from smoothed motor unit spike trains using the MATLAB scripts, and recruitment and derecruitment thresholds were determined by obtaining the force at the instant of the first and last spikes, respectively, for each motor unit. Ascending duration was defined as the interval during which the motor unit maintained continuous discharge from recruitment to peak force, while descending duration captured the interval from peak to derecruitment. From these discharge durations, we computed the self-sustained duration (SSD; Afsharipour et al., 2020; Mohammadalinejad et al., 2024), which is the relative difference in time spent on the descending phase of the ramp compared to the ascending phase, normalized to the entire duration of discharge (i.e., in percentage).

#### Motor unit tracking across conditions

To identify the same motor units across different conditions (CONTROL, TAP, LOW, HIGH), motor unit filters previously identified by the CKC method during each condition were applied to the HDsEMG signals recorded at other conditions at their respective intensities. To achieve this, the arm cycling trials were first edited, and then were concatenated with the other conditions of the same contraction intensity. Motor unit filters from each individual condition were then applied to the concatenated recordings, generating motor unit spike trains across all conditions for a given contraction intensity. DEMUSE identified duplicates when a motor unit spike train shared a 33% or greater match with other motor unit spike trains. After removing duplicate motor units detected during multiple conditions, motor unit spike trains were once again manually inspected and edited.

#### Paired motor unit analysis

A common metric used to estimate the contribution of PICs to motoneuron discharge in humans is delta frequency (ΔF), which quantifies the amount of onset-offset hysteresis of a higher-threshold motor unit with respect to the discharge rate of a lower threshold motor unit. ΔF for a given motor unit (test unit) is quantified as the change in discharge rate of a lower threshold motor unit (reporter unit) between the recruitment and derecruitment instance of the test unit (Gorassini et al., 2002a). To account for the possibility of each test unit pairing with multiple lower-threshold reporter units, ΔF was calculated as a unit-wise value as first presented by Hassan et al. (2020), representing the average change in discharge rate across all valid reporter-test pairs for each test unit. Motor unit pairs were required to meet criteria to be included in ΔF calculations: 1) the test motor unit had to be recruited at least 1 second after the reporter motor unit to allow sufficient time for full PIC activation (Bennett et al., 2001; Hassan et al., 2020; Powers et al., 2008), 2) test unit-reporter unit pair exhibited rate-rate correlations of r^2^ 0.7, ensuring that motor unit pairs likely received common synaptic drive (Gorassini et al., 2002b; Udina et al., 2010; Wilson et al., 2015) and 3) the reporter unit modulated its discharge by at least 0.5 pps while the test unit was active (Stephenson et al., 2011). Simulations suggest that ΔF is sensitive to changes in neuromodulation, as well as the amount and pattern of inhibition (Beauchamp et al., 2023). An example of this method is displayed in Figure 3.

**Figure 3.**
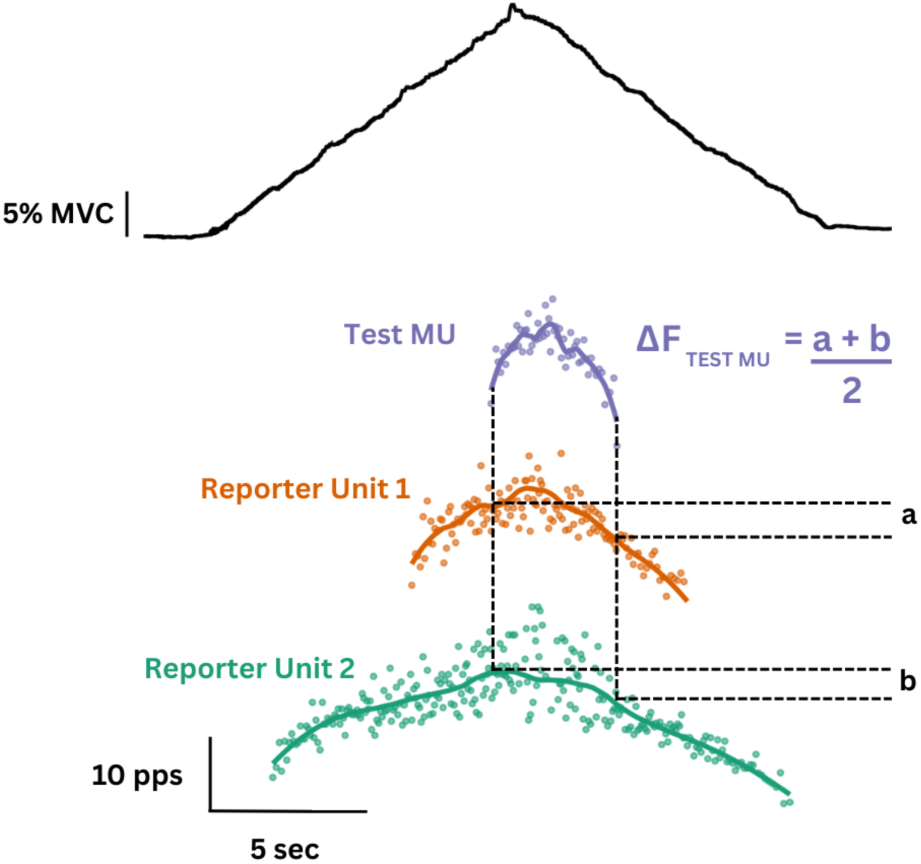
The paired motor unit analysis technique. This technique estimates the contribution of PICs to motoneuron discharge in humans by quantifying the difference between the discharge rate of a lower-threshold (control unit) motor unit at the recruitment and de-recruitment of a higher-(test) motor unit, which measures the hysteresis in the discharge rate (Gorassini et al, 2002). In this case, the ΔF is the average value obtained from the ΔF of the purple test unit with respect to the discharge rate of the orange and green reporter units.

#### Geometric Analysis

Another metric, known as brace height (BH) was used to quantify the non-linearity of the ascending portion of the motor unit discharge rates with respect to force output (Beauchamp et al., 2023). Simulations have demonstrated BH may provide greater insight into the role of neuromodulatory inputs on motor unit discharge, as this metric is less sensitive to changes in the pattern of inhibition (Beauchamp et al., 2023). BH was calculated as the maximum orthogonal deviation between the smoothed discharge rate and an expected linear increase from motor unit recruitment to peak discharge. The maximal deviation was defined as ‘brace height’. To account for scaling effects due to discharge range, BH was normalized to the height of a right triangle formed between the recruitment and peak discharge. Normalization of BH allows for comparisons across motor units and reflects the relative degree of PIC-mediated amplification in discharge rate.

### Statistical Procedures

Statistical analyses were performed using R Statistical Software (v4.5.0; R Core Team 2025). To determine if variables of interest were predicted by the fixed effects of condition, intensity, and their interactions, we used linear mixed-effects models with covariates of recruitment threshold (lmer R package, v1.1.27.1 (Bates et al., 2015)). Because motor units were tracked across conditions, the motor unit identifier was nested within participant and included as a random intercept. Model statistics are reported as chi-squared and degrees of freedom. To determine significance, we applied Satterthwaite’s method for degrees of freedom (lmerTest R package; v3.1.3;(Kuznetsova et al., 2017)). Results are reported as estimated marginal means (emmeans R package, v1.8.0;(Lenth et al., 2025) ± 95% confidence limits. Effect sizes (Cohen’s d) were calculated from the estimated marginal means to determine the standardized magnitude of the effect of condition within the low and high intensity contractions compared to control from the model. All data were visualized in R (ggplot R package, v3.3.6; (Wickham et al., 2025).

## RESULTS

In this study, we examined differences in motor unit discharge characteristics between conditions during isometric triangular shaped contraction of 25% and 50% MVC. Data were only stratified by sex when characterizing the motor units identified. For females, we decomposed a total of 30 motor unit spike trains at 25% and 17 motor unit spike trains at 50% that could be tracked across all four conditions. Seven female participants were recruited for the study, but due to poor decomposition or task performance, three female’s data were removed due to poor signal quality. For males, we decomposed a total of 128 motor unit spike trains at 25%, and 97 motor unit spikes trains were decomposed at 50% that could be tracked across all four conditions. Out of the ten males that participated, one male’s data was removed due to poor signal quality and one due to poor task performance. Motor units that could not be tracked across all four conditions were not used for further analysis. *Table 1* shows the number of units identified and tracked for each participant. Figure 4 show examples of tracked motor units across conditions during 50% MVC contractions

**Figure 4.**
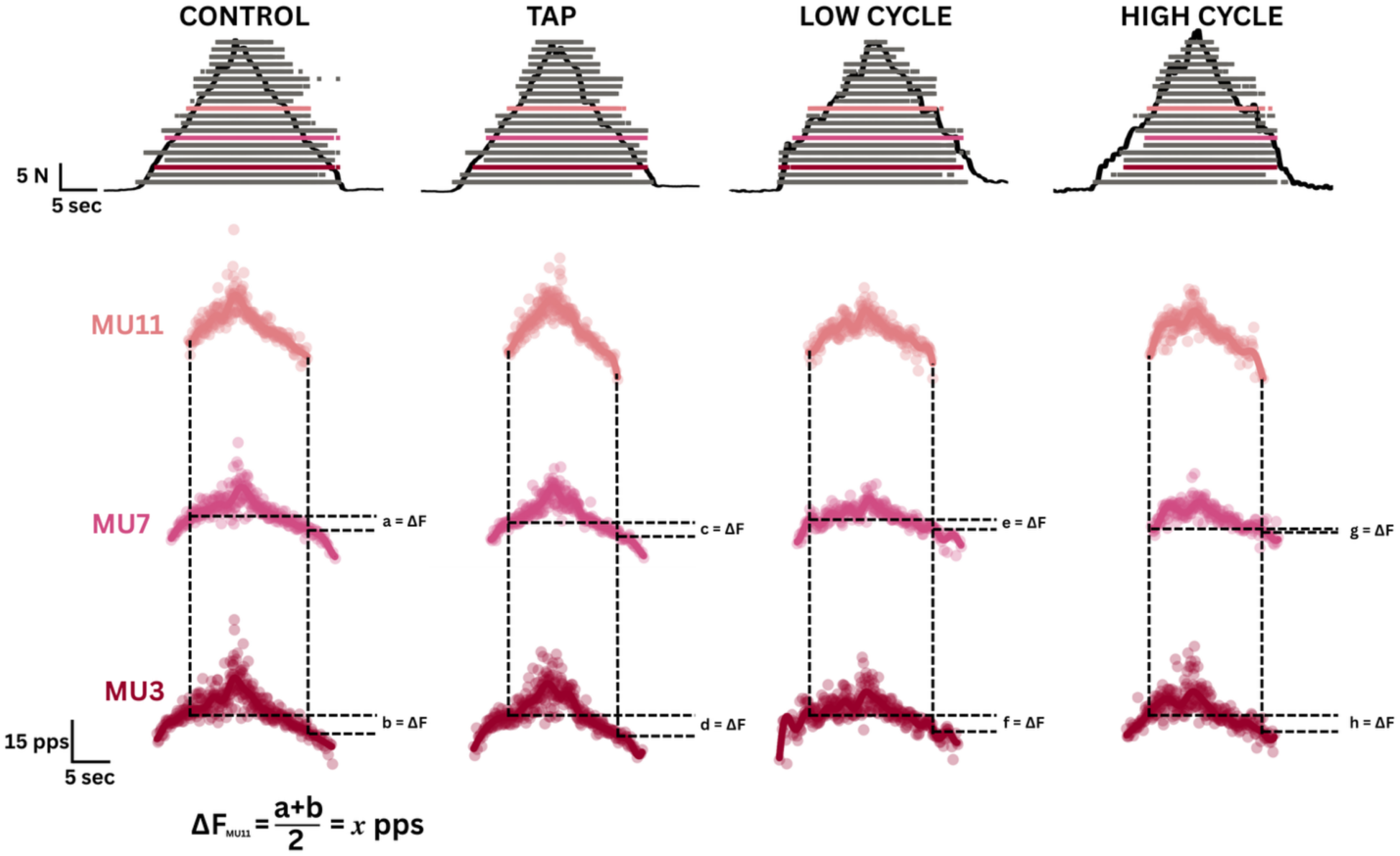
Example data. A single participant’s data during the CONTROL, TAP, LOW, and HIGH conditions during a 50% MVC contraction. The top of the figure depicts the force trace in the thick dark grey trace that was produced by the participant performing a dorsiffexion contraction. Overlayed on the force trace is a raster plot that show motor unit discharges and the order in which they were recruited, with red units recruited at a low force output and blue units recruited at higher forces. Below the force trace and raster plot are the smoothed discharge rates of three identical motor units. This figure displays how the paired motor unit analysis is performed to calculate ΔF.

**Table 1.**
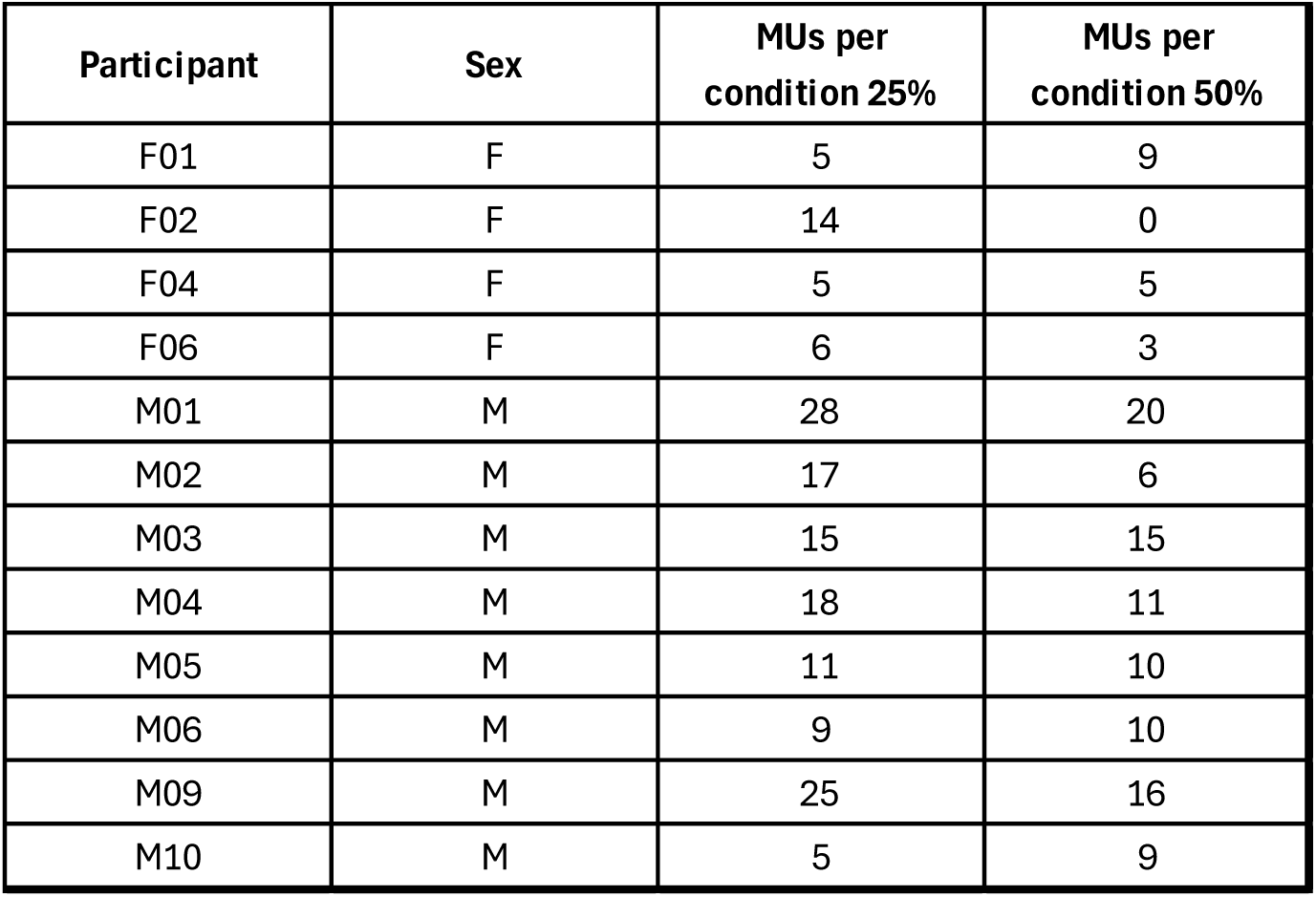
Motor units spike trains decomposed for each participant across intensities.

### Motor unit recruitment and derecruitment threshold distributions are similar across conditions

Motor unit recruitment was compared across four conditions (CONTROL, TAP, LOW CYCLE, HIGH CYCLE) at both 25% and 50% MVC using linear mixed-effects models with random intercepts for motor units nested within participants. A likelihood ratio test indicated that adding the Condition × Intensity interaction did not improve model fit (χ²(3) = 0.97, p = 0.81); therefore, the interaction model was not used. This model revealed a robust main effect of intensity, with recruitment thresholds approximately 16% MVC higher at 50% compared to 25% MVC (p < 0.00001). In contrast, there was no significant main effect of condition (all Tukey-adjusted p ≥ 0.49). Estimated marginal means were highly consistent across conditions, ranging from 18.7 ± 0.67% MVC (LOW) to 19.0 ± 0.67% MVC (CONTROL), as shown in Figure 5. Descriptively, recruitment thresholds spanned 10.7–11.1% MVC at 25% MVC and 26.6–27.0% MVC at 50% MVC. These results indicate that motor unit recruitment remained consistent across contraction conditions, regardless of contraction intensity.

**Figure 5.**
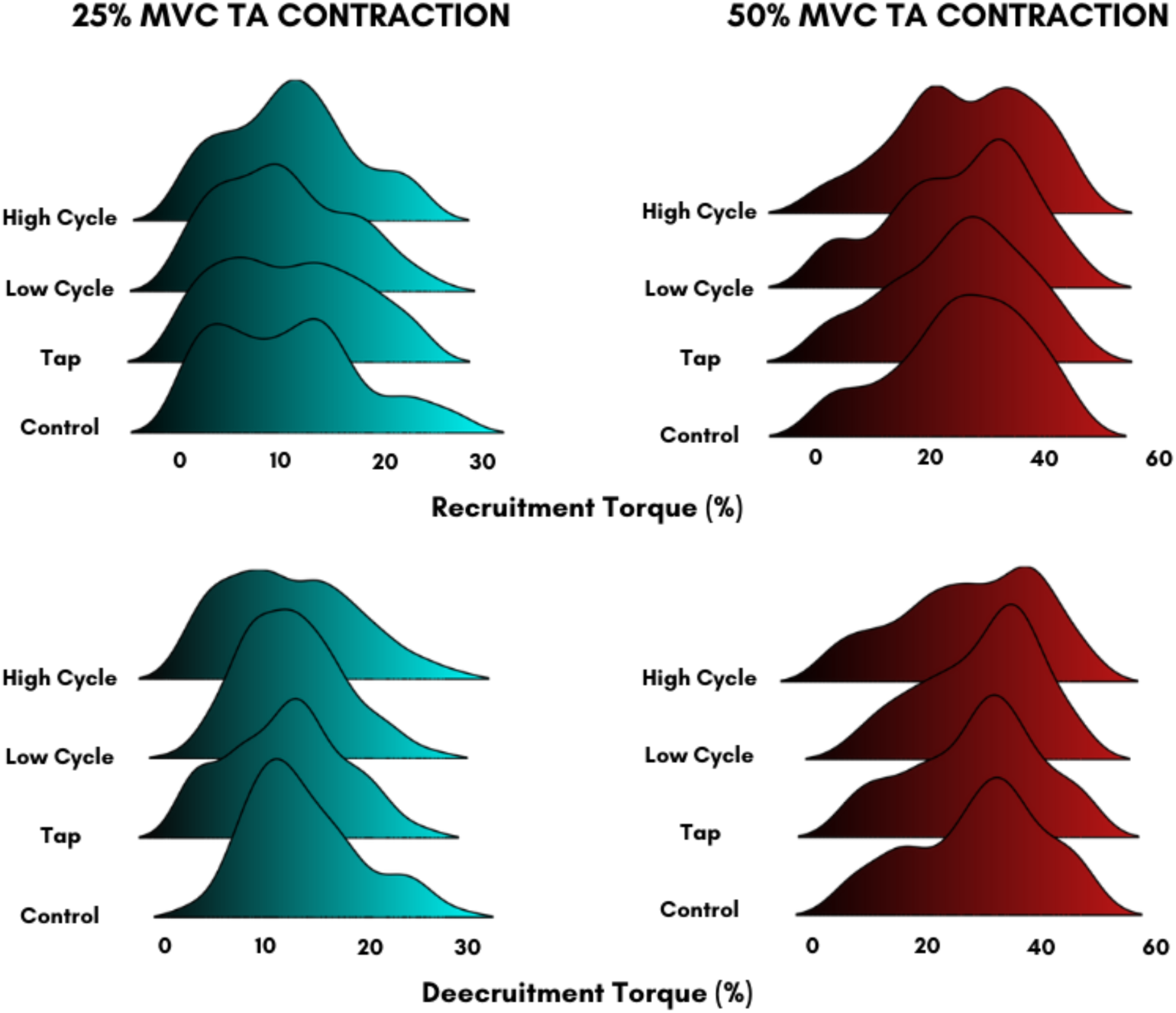
Distribution of recruitment and derecruitment thresholds of motor units throughout intensities and across conditions. Motor units recruited at 25% MVC are shown in blue, and motor units recruited at 50% are shown in red. For each figure, the x-axis shows the percentage of maximal voluntary contraction (MVC). The y-axis indicates probability density, not raw counts, which shows the relative concentration of recruitment thresholds in each condition.

**Table 2.**
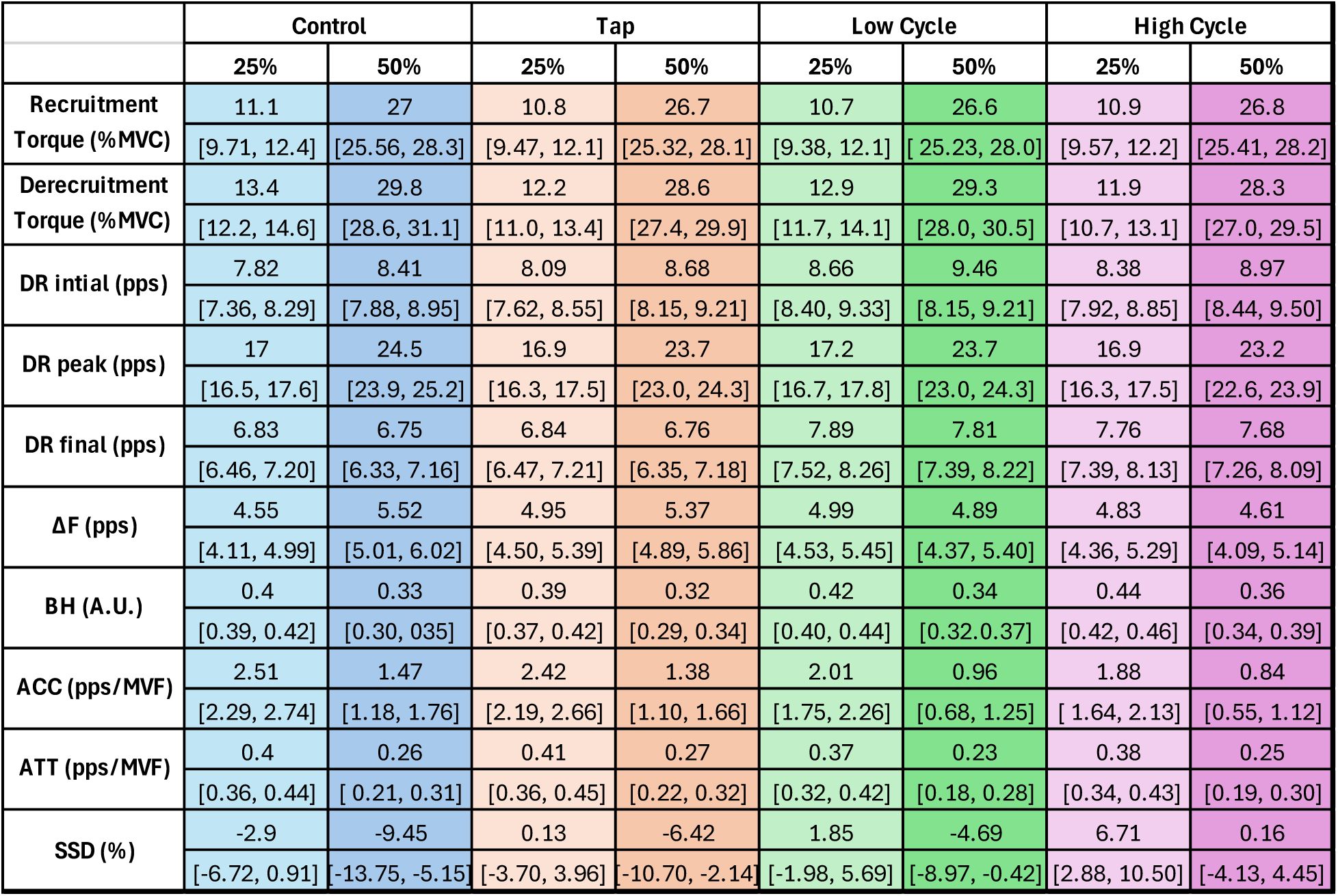
Estimated marginal means (95% CI) for all motor unit variables.

### ΔF declines during high-intensity contractions accompanied by arm cycling

Estimates of PIC-induced hysteresis were analyzed across conditions and intensities and fell within the normal range of values reported in previous studies (Afsharipour et al., 2020; Gomes et al., 2024; Jenz et al., 2023; G. E. Pearcey et al., 2022; Škarabot, Beauchamp, et al., 2025). A significant interaction between condition and intensity was detected [χ²(3) = 13.44, P = 0.0038]. At 25% MVC, there were no significant differences in ΔF between conditions (all p > 0.26), with an overall mean of 4.83 ± 0.23 pps across conditions as seen in Figure 6, but at 50% MVC, ΔF was significantly lower in the HIGH compared to CONTROL (P = 0.0048, d = 0.51) and TAP (P = 0.0213). Although pairwise comparisons were not significant, ΔF tended to be lower in the LOW condition compared to CONTROL (P = 0.0908). These findings indicate that, while estimates of the contribution of PICs to motoneuron discharge generally increase with contraction intensity, this effect is dampened in the presence of higher cadence upper body rhythmic motor output.

**Figure 6.**
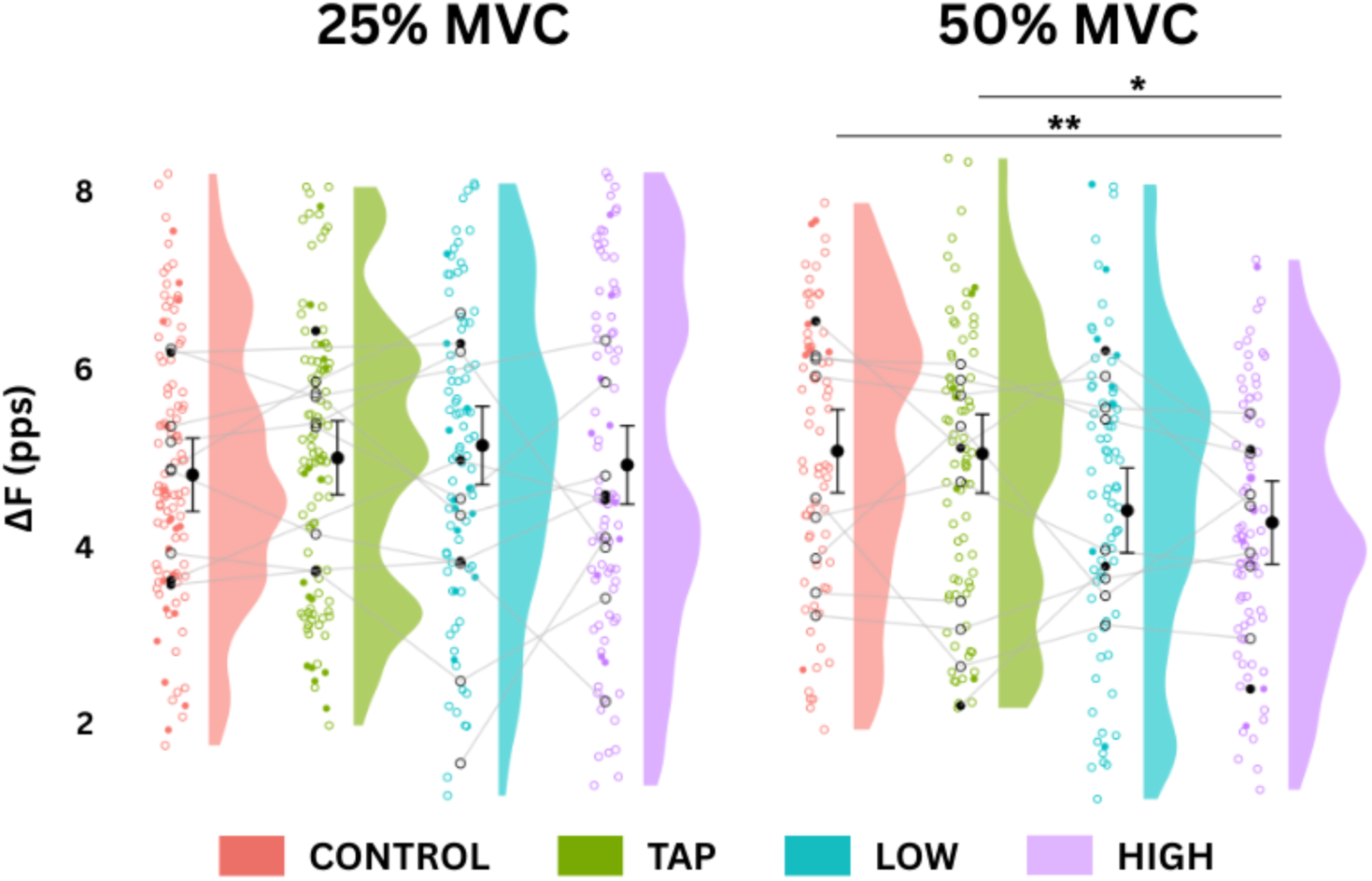
Changes in ΔF across conditions within intensities. The data is presented by intensity across conditions is represented with a half-violin (density) plot, overlaid with raw participant-level data-colored **open** circles **(male)** and **closed** circles **(female)** and lines connecting individual participants’ mean values across conditions (gray lines). Estimated marginal means (EMMs) with 95% confidence intervals are plotted as larger solid black circles with error bars, slightly nudged to the right of the distribution (*P < 0.05, **P < 0.01).

### Non-linearity of ascending discharge rates is increased during arm cycling

To assess the non-linearity of the ascending motor unit discharge rate, a quasi-geometric analysis of the individual motor units discharge rates with respect to force output was used. As shown in Figure 7, we found that BH was significantly influenced by both contraction intensity and condition, but not their interaction [χ²(3) = 2.00, P = 0.572]. Specifically, BH was lower during contractions to 50% compared to 25% MVC across all conditions [χ²(1) = 35.01, p < 0.000001, d = 0.503], consistent with reduced motoneuron discharge non-linearity at higher contraction intensities (Škarabot et al., 2025). Across intensities, condition exerted a small but significant effect [χ²(3) = 14.28, P = 0.0026], with values being highest in the HIGH condition (0.439 ± 0.0106), followed by LOW (0.418 ± 0.0111), CONTROL (0.404 ± 0.0098), and TAP (0.394 ± 0.0103). Post hoc comparisons revealed that BH in the HIGH condition was significantly larger than CONTROL (P = 0.0256; d = 0.117) and TAP (P = 0.0023; d = 0.148), indicating a subtle yet consistent elevation in BH during repetitive locomotor actions.

**Figure 7.**
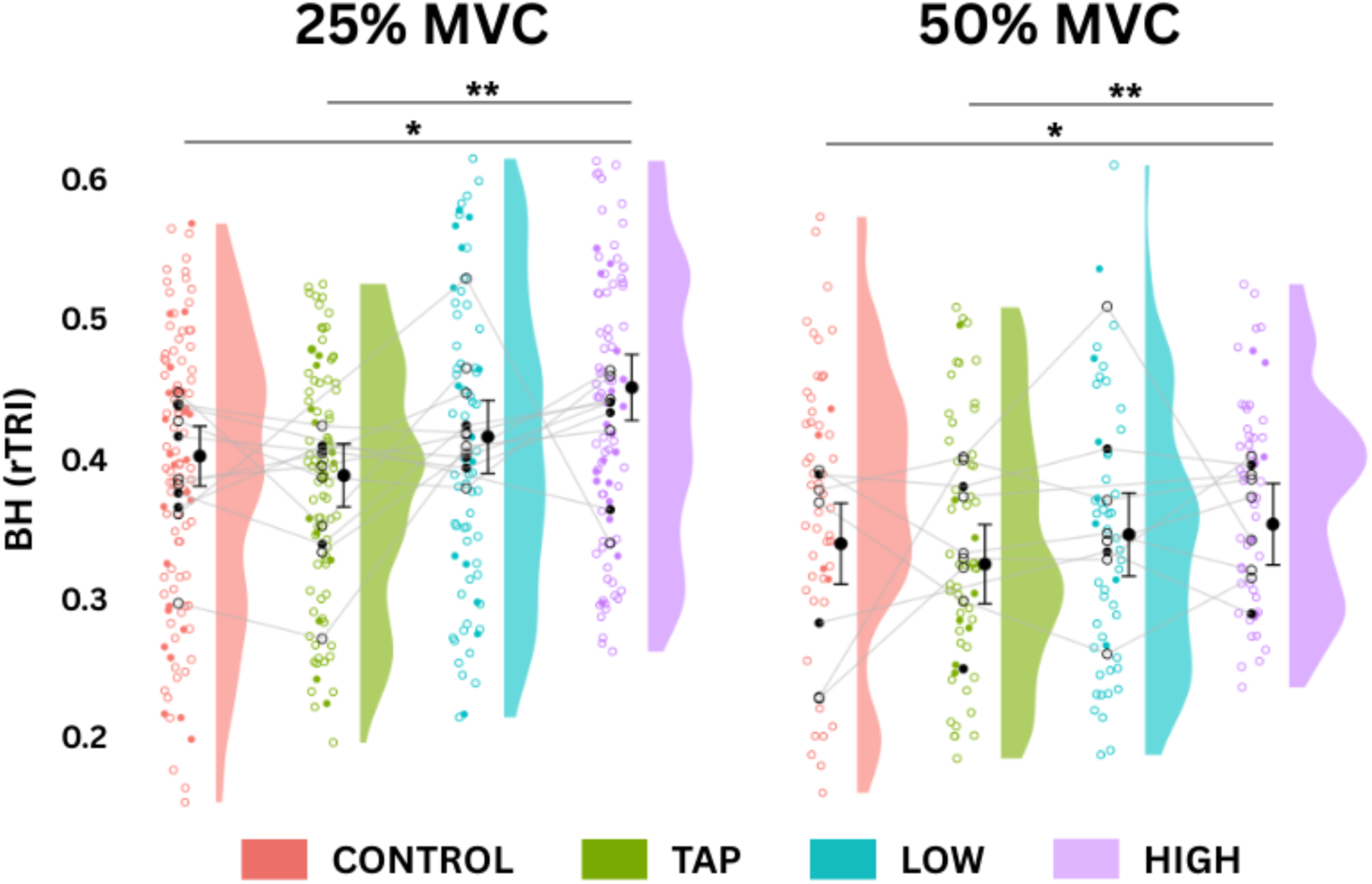
Changes in BH across conditions within intensities. The data is presented by intensity across conditions is represented with a half-violin (density) plot, overlaid with raw participant-level data-colored **open** circles **(male)** and **closed** circles **(female)** and lines connecting individual participants’ mean values across conditions (gray lines). Estimated marginal means (EMMs) with 95% confidence intervals are plotted as larger solid black circles with error bars, slightly nudged to the right of the distribution (*P < 0.05, **P < 0.01).

Acceleration (ACC) slope (i.e., secondary discharge range slope), as shown in Figure 8, displayed main effects of condition [χ²(3) = 26.89, P < 0.000001] and contraction intensity [χ²(1) = 29.88, P < 0.000001], with no significant interaction [χ²(3) = 1.26, P = 0.739]. Across intensities, ACC was significantly lower during 50% MVC contractions compared to 25% MVC (d = 0.544). Across intensities, ACC was significantly reduced in both LOW (P = 0.0030; d = 0.147) and HIGH (P = 0.0001; d = 0.186) relative to CONTROL, and HIGH was also significantly lower than TAP (P = 0.0011; d = 0.158). These differences suggest that rhythmic upper body tasks attenuate the acceleration in motoneuron discharge just after recruitment, particularly in more demanding cycling conditions.

**Figure 8.**
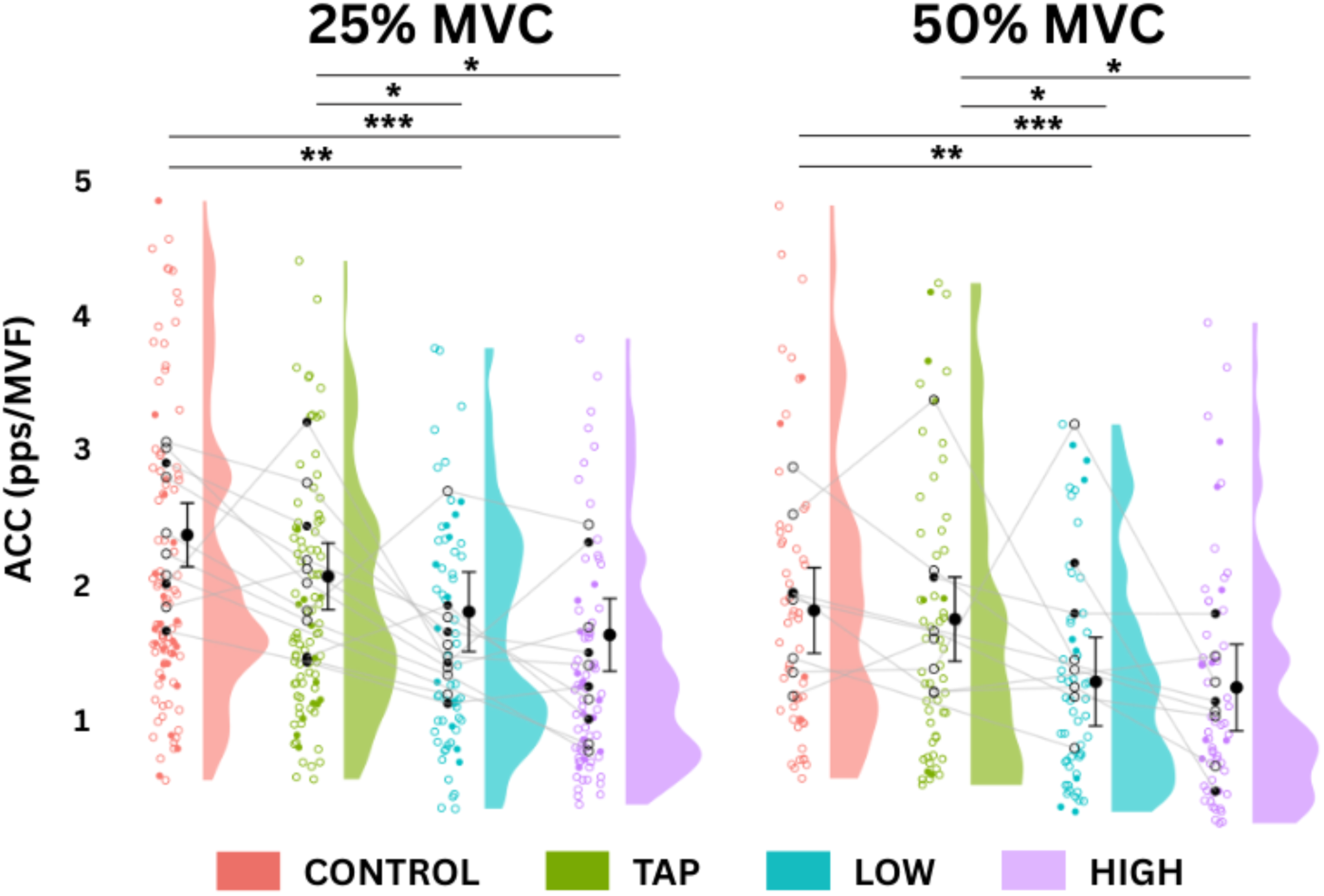
Changes in ACC across conditions within intensities. The data is presented by intensity across conditions is represented with a half-violin (density) plot, overlaid with raw participant-level data-colored **open** circles **(male)** and **closed** circles **(female)** and lines connecting individual participants’ mean values across conditions (gray lines). Estimated marginal means (EMMs) with 95% confidence intervals are plotted as larger solid black circles with error bars, slightly nudged to the right of the distribution (*P < 0.05, **P < 0.01, ***P< 0.001).

Attenuation (ATT) slope (i.e., tertiary discharge range slope) was significantly affected by contraction intensity [χ²(1) = 25.44, P < 0.000001, d = 0.503], with values being lower during 50% MVC contractions compared to 25% MVC (Figure 9). No significant main effect of condition [χ²(3) = 2.28, P = 0.516] or condition-by-intensity interaction [χ²(3) = 1.18, P = 0.757] was detected. Thus, ATT did not differ significantly between conditions.

**Figure 9.**
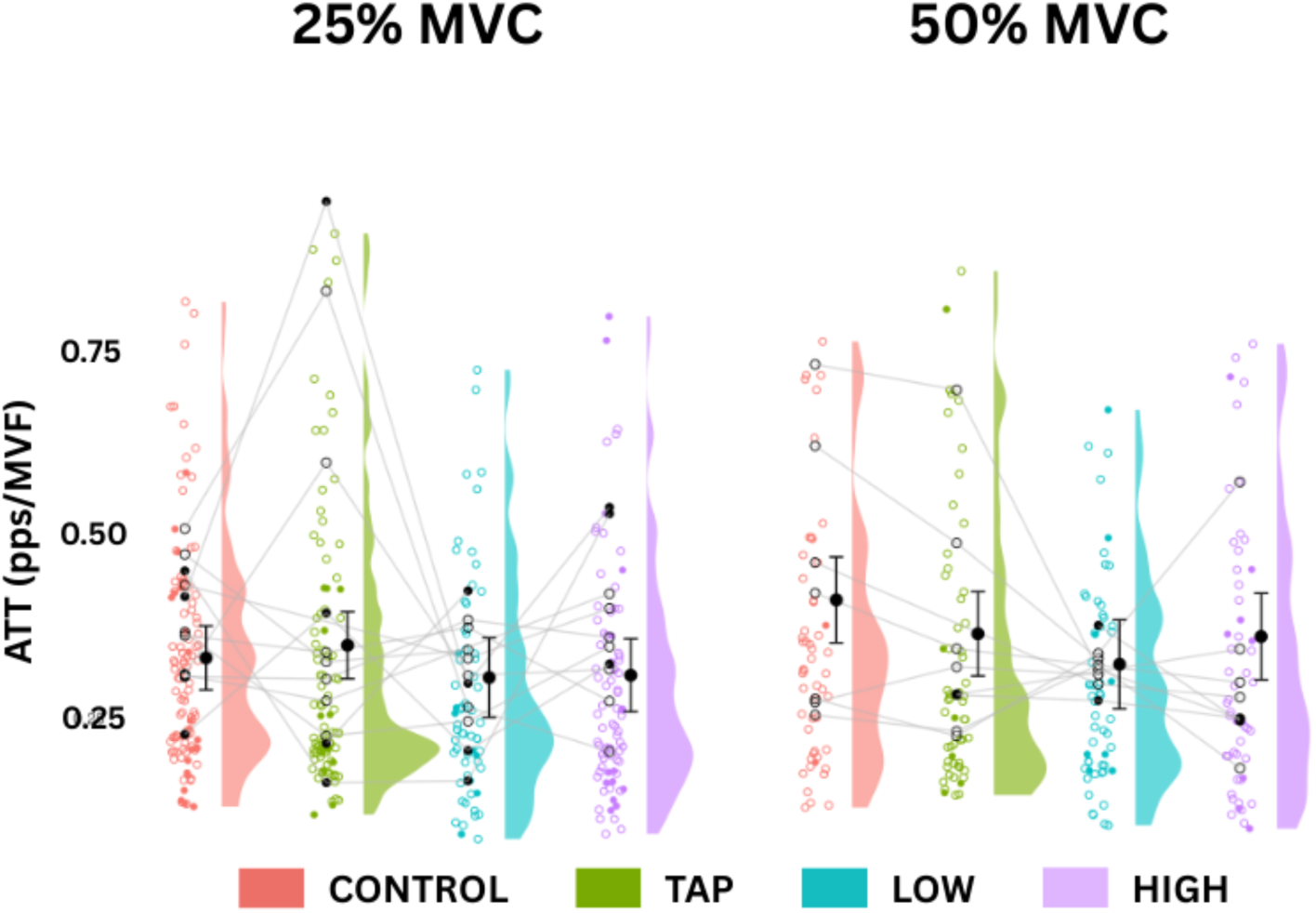
Changes in ATT across conditions within intensities. The data is presented by intensity across conditions is represented with a half-violin (density) plot, overlaid with raw participant-level data-colored **open** circles **(male)** and **closed** circles **(female)** and lines connecting individual participants’ mean values across conditions (gray lines). Estimated marginal means (EMMs) with 95% confidence intervals are plotted as larger solid black circles with error bars, slightly nudged to the right of the distribution.

For self-sustained duration (SSD), there were significant main effects of intensity [χ²(1) = 4.12, P = 0.042] and condition [χ²(3) = 15.00, P = 0.002], but not their interaction. SSD was less at 50% MVC compared to 25% MVC (d = 0.101) and increased (less negative) in HIGH relative to CONTROL (P < 0.0001; d = 0.205) and TAP (P = 0.0001; d = 0.140) as seen in Figure 10. LOW also exhibited more SSD compared to CONTROL (P = 0.0119; d = 0.101) and LOW had significantly less SSD than HIGH (P = 0.0098; d = 0.103). This suggests that there was more prolonged discharge during cycling conditions relative to CONTROL and TAP.

**Figure 10.**
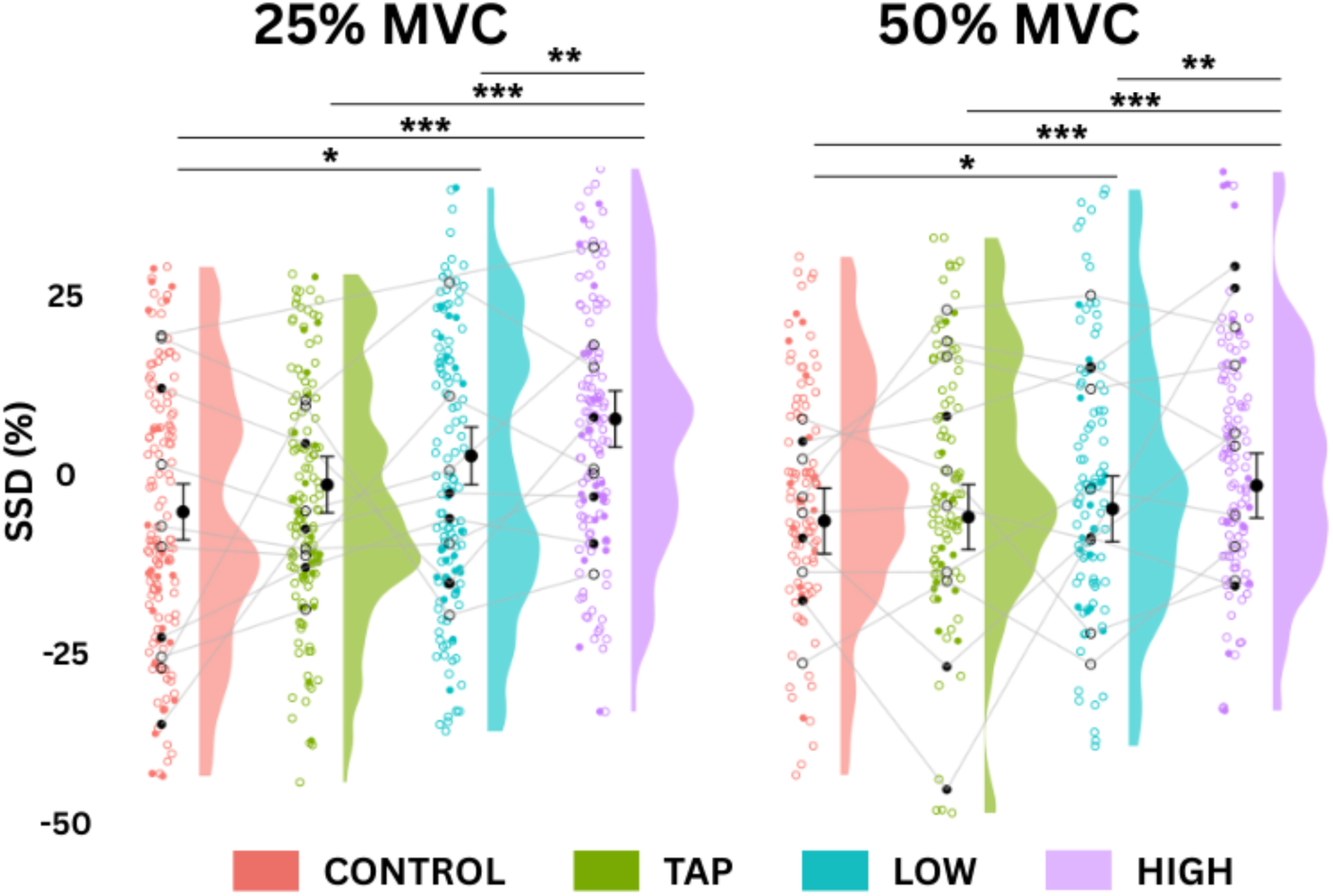
Changes in SSD across conditions within intensities. The data is presented by intensity across conditions is represented with a half-violin (density) plot, overlaid with raw participant-level data-colored **open** circles **(male)** and **closed** circles **(female)** and lines connecting individual participants’ mean values across conditions (gray lines). Estimated marginal means (EMMs) with 95% confidence intervals are plotted as larger solid black circles with error bars, slightly nudged to the right of the distribution (*P < 0.05, **P < 0.01, ***P<0.001).

## DISCUSSION

The objective of this study was to investigate the effects of locomotor output of the arms on the excitability of motoneurons in the stationary non-exercising legs. Results indicate that there were reductions in ΔF during ramp contractions to 50% accompanied by arm cycling, which is directly contrary to our hypothesis. Further analysis of discharge rate parameters analyzed in conjunction with the reduced ΔF helped us glean further insight into the changes induced by remote locomotor activity. In particular, the increased non-linearity of ascending discharge rates may indicate enhanced neuromodulation (i.e., 5-HT facilitation of motoneuron discharge), whilst the reduction in ΔF may be the result of more proportional or increased levels of inhibition, which we elaborate on below.

Upper body locomotor output did not increase our main estimate of PIC magnitude (ΔF) in the TA muscle. At low contraction intensities ΔF remained unchanged during arm cycling, whereas during high contraction intensities, ΔF decreased during the arm cycling tasks. The lack of increase in ΔF was initially surprising, as we expected monoamine concentrations to increase with locomotor behaviour and thus proportionally increase voltage-gated PIC channels (Heckman et al., 2009); however, the decrease in ΔF may be explained by PIC’s sensitivity to inhibitory inputs. There is evidence in animal models by which disynaptic Ia reciprocal inhibition can reduce PIC activity (Hyngstrom et al., 2007; Kuo et al., 2003). Similarly, there is preliminary evidence in humans that local inhibitory inputs can reduce ΔF in humans (Mesquita et al., 2022; Pearcey et al., 2022). The suppression of ΔF in the leg during arm cycling may have been a consequence of an inhibition from spinal circuits during locomotor behaviour. While the leg is stationary, rhythmic arm cycling has been shown to reduce H-reflex amplitude (Loadman & Zehr, 2007), suggesting that presynaptic inhibition increases in the leg during locomotor output (Frigon et al., 2004; Pearcey & Zehr, 2019)

There is limited research exploring how upper body activity affects PICs in the lower limb. An exception is a study by Orssatto et al. (2022), which had participants perform remote handgrip contractions prior to isometric ramp contractions in the leg. They reported that these remote efforts facilitated estimates of the contribution of PICs to motoneuron discharge, as quantified by ΔF. In contrast, our findings suggest a different pattern, reflecting the involvement of distinct neural pathways. Specifically, rhythmic locomotor activity engages CPGs that provide a unique balance of excitatory and inhibitory input to spinal motoneurons, differing from the input produced by static upper limb contractions (Frigon et al., 2004; Loadman & Zehr, 2007; Pearcey & Zehr, 2019; Zehr et al., 2004).

Although PICs are sensitive to inhibition, and as such were reflected in our ΔF measures, there still may have been an increase in neuromodulation, via monoaminergic inputs (Binder et al., 2020; Heckman et al., 2009). One the main sources of neuromodulatory input is 5-HT released from Raphe neurons in the brainstem (Johnson et al., 2017). Locomotor activity increases the discharge rate of descending serotonergic neurons, which release 5-HT onto motoneurons throughout the spinal cord; this increase is linked to walking speed in cats (Jacobs et al., 2002). In a study by Basile et al., they assessed motor unit discharge rates recorded from intramuscular EMG during arm cycling. They suggested that CPG activity may have resulted in greater activation of PICs, which may have contributed to the increased motor unit discharge rates during cycling compared to the intensity-matched isometric contractions. Given the highly diffuse nature of 5-HT, it is likely involved in mediating state changes of motoneuron excitability (Basile et al., 2025; MacDonell et al., 2015; Power et al., 2010). Because locomotor activity is associated with 5-HT release in cats (Jacobs et al., 2002), and despite the precise mechanisms underlying increased motoneuron excitability during arm cycling in humans remaining unclear (Basile et al., 2025), it is highly likely that some degree of neuromodulation occurred during the arm cycling trials in our study, as indicated by the increase in non-linearity and SSD, especially during higher cadence arm cycling.

Despite the observed decreases in ΔF observed during rhythmic activity in the upper body, BH increased. Unlike ΔF, which is sensitive to both patterns of inhibition and neuromodulation, BH is mainly sensitive to changes in neuromodulation (Beauchamp et al., 2023), which allows us to disentangle locomotor effects on motoneuron discharge rate patterns. Based on our findings, upper body locomotion increases BH during low and high intensity contractions in the leg; this effect is slightly dampened during the higher intensity contractions. The increase in BH is consistent with existing evidence showing that activation of the Raphe nuclei increased during locomotor activity, as demonstrated in cat models (Jacobs et al., 2002).

Although the overall reduction in ΔF during higher intensity contractions, on the other hand, may also reflect a “spillover effect” of the release of 5-HT, this trend contradicts with previous findings showing that ΔF measured across a range of intensities during isometric contractions tends to increase as contraction level increases (Beauchamp et al., 2023; Škarabot et al., 2025; Škarabot et al., 2025). Although the inhibitory effects of 5-HT are less frequently observed, preparations utilizing adult turtle spinal cords have shown variable effects of 5-HT on motoneuron excitability. Cotel et al. (2013) reported that ‘moderate’ release of 5-HT release activated 5-HT2 receptors and motoneuron excitability increased; however, as 5-HT release increased and passed a certain threshold, 5-HT₁_A_ receptors were activated, leading to a reduction of motoneuron excitability. Increased activation of serotonergic fibers may release enough 5-HT to saturate 5-HT₂ receptors, leading to spillover and subsequent activation of 5-HT₁_A_ receptors. An in vivo human study showed that elevated extracellular levels of endogenously released 5-HT impaired maximal torque production, suggesting that 5-HT release onto motoneurons may contribute to central fatigue (Kavanagh et al., 2019). In our current study we were unable to monitor whether a spillover of 5-HT onto 5-HT₁_A_ receptors occurred; however, the combination of arm cycling and higher intensity TA contractions may have generated sufficient serotonergic drive to shift the net neuromodulatory effects of 5-HT from facilitating to slightly suppressing motoneuron excitability compared to lower intensity TA contractions.

### Methodological Considerations

A key concern was ensuring that any observed changes weren’t simply due to cognitive demands of dual-tasking and incidental upper-body movement, which could increase descending voluntary drive to a level at which it could alter estimates of the contribution of PICs to motoneuron discharge. To address this, we introduced the TAP condition, which involved tapping with the right index finger cued by auditory beeps during dorsiflexion ramp contractions. This task served to test whether a mild cognitive challenge combined with subtle distal upper-limb movement could modulate motoneuron excitability. In all measures the TAP condition did not differ from CONTROL, which instills confidence in our results with respect to our LOW and HIGH cycling conditions. This suggests that changes seen in the cycling condition were more tightly coupled the unique aspects of locomotor output, rather than cognitive demand of dual tasking.

Following the target force trace while simultaneously performing the arm cycling task proved challenging for participants. During the arm cycling conditions, some participants experienced torso movement, which introduced additional difficulty. The minor sway of the torso caused slight fluctuations in force output, complicating their performance in producing smooth, linear increases and decreases in force during the ascending and descending phases. This variability was also reflected in the decomposition of the HDsEMG signals, where certain motor unit spike trains exhibited patterns mirroring the force fluctuations. In some cases, these fluctuations may have contributed to early derecruitment of motor units, particularly during down ramp phase of the TA contractions. The exclusion of participants who exhibited excessive torso movements and fluctuations in force and the inclusion of the TAP control condition confirmed that these changes were specific to locomotor output rather than cognitive effort or incidental movement effects, underscoring the unique capacity of rhythmic arm activity to modulate motoneuron behavior in the lower limb.

A further consideration relates to the intensity of the arm cycling task. In the present study, wattage was measured using an analogue scale integrated into the arm cycle ergometer, and all participants, regardless of sex or training status, cycled at the same absolute intensity. A more precise approach to standardizing workload would involve determining each participant’s maximal arm cycling capacity (e.g., through an incremental workload test) and then setting the experimental load at a fixed percentage of this maximum. This method would ensure that all participants cycled at an equivalent relative intensity, potentially reducing inter-individual variability in task difficulty and physiological response.

Finally, in this study, we only included motor units that could be tracked across conditions within a single intensity. Although this approach controlled for unit-specific properties, helped to eliminate inter-unit variability as a confounding factor, isolate condition effects, and allowed the comparison of motor units receiving the same synaptic input (Orssatto et al., 2021; Trajano et al., 2020), we may have missed effects that manifest from recruitment of unique units in one condition compared to another.

## CONCLUSION

This study examined the influence of rhythmic upper body locomotor activity on motoneuron excitability innervating the stationary, non-exercising tibialis anterior muscle during low- and high-intensity isometric contractions. While recruitment and derecruitment thresholds remained unchanged across conditions, upper body cycling at higher cadences reduced ΔF, suggesting that the amount or pattern of inhibitory input to motoneurons is altered during remote locomotor activity, potentially via spinal circuits. In contrast, non-linearity and self-sustained discharge increased across cycling conditions, particularly at lower contraction intensities, which indicates an increase in neuromodulatory drive, likely mediated by 5-HT. These findings suggest that a complex interplay between excitation and inhibition at the spinal level during rhythmic upper limb activity.

## Notes

### Competing Interest Statement

The authors have declared no competing interest.

### Funding

- Natural Sciences and Engineering Research Council: Discovery Grant [2023-05862]
- Azrieli Foundation: Collaboration on Motor Planning, Execution and Resilience Network Grant

## References

Afsharipour, B., Manzur, N., Duchcherer, J., Fenrich, K. F., Thompson, C. K., Negro, F., Quinlan, K. A., Bennett, D. J., & Gorassini, M. A. (2020). Estimation of self-sustained activity produced by persistent inward currents using firing rate profiles of multiple motor units in humans. Journal of Neurophysiology, 124(1), 63–85.

Alahmari, S. K., Orssatto, L. B. R., Shield, A. J., & Trajano, G. S. (2026). Effects of arm-cycling exercise during triceps surae neuromuscular electrical stimulation on torque output and fatigue. European Journal of Applied Physiology, 126(1), 95–105. 10.1007/s00421-025-05879-y

Basile, D. C., Wira, A. D., Rice, C. L., & Power, K. E. (2025). Investigating motor unit firing rates during arm cycling compared with intensity-matched isometric contractions in humans. Journal of Neurophysiology, 134(1), 162–170. 10.1152/jn.00128.2025

Bates, D., Mächler, M., Bolker, B., & Walker, S. (2015). Fitting Linear Mixed-Effects Models Using lme4. Journal of Statistical Software, 67, 1–48. 10.18637/jss.v067.i01

Beauchamp, J. A., Khurram, O. U., Dewald, J. P. A., Heckman, C. J., & Pearcey, G. E. P. (2022). A computational approach for generating continuous estimates of motor unit discharge rates and visualizing population discharge characteristics. Journal of Neural Engineering, 19(1). 10.1088/1741-2552/ac4594

Beauchamp, J. A., Pearcey, G. E. P., Khurram, O. U., Chardon, M., Wang, Y. C., Powers, R. K., Dewald, J. P. A., & Heckman, C. J. (2023). A geometric approach to quantifying the neuromodulatory effects of persistent inward currents on individual motor unit discharge patterns. Journal of Neural Engineering, 20(1), 016034. 10.1088/1741-2552/acb1d7

Bennett, D. J., Li, Y., & Siu, M. (2001). Plateau potentials in sacrocaudal motoneurons of chronic spinal rats, recorded in vitro. Journal of Neurophysiology, 86(4), 1955–1971.

Binder, M. D., Powers, R. K., & Heckman, C. J. (2020). Nonlinear input-output functions of motoneurons. Physiology, 35(1), 31–39.

Borzuola, R., Nuccio, S., Scalia, M., Parrella, M., Del Vecchio, A., Bazzucchi, I., Felici, F., & Macaluso, A. (2023). Adjustments in the motor unit discharge behavior following neuromuscular electrical stimulation compared to voluntary contractions. Frontiers in Physiology, 14. 10.3389/fphys.2023.1212453

Cotel, F., Exley, R., Cragg, S. J., & Perrier, J.-F. (2013). Serotonin spillover onto the axon initial segment of motoneurons induces central fatigue by inhibiting action potential initiation. Proceedings of the National Academy of Sciences, 110(12), 4774–4779. 10.1073/pnas.1216150110

Dragert, K., & Zehr, E. P. (2009). Rhythmic arm cycling modulates Hoffmann reflex excitability differentially in the ankle flexor and extensor muscles. Neuroscience Letters, 450(3), 235–238. 10.1016/j.neulet.2008.11.034

Feraboli-Lohnherr, D., Barthe, J.-Y., & Orsal, D. (1999). Serotonin-induced activation of the network for locomotion in adult spinal rats. Journal of Neuroscience Research, 55(1), 87–98. 10.1002/(SICI)1097-4547(19990101)55:1<87::AID-JNR10>3.0.CO;2-#

Frigon, A. (2017). The neural control of interlimb coordination during mammalian locomotion. Journal of Neurophysiology, 117(6), 2224–2241. 10.1152/jn.00978.2016

Frigon, A., Collins, D. F., & Zehr, E. P. (2004). Effect of Rhythmic Arm Movement on Reflexes in the Legs: Modulation of Soleus H-Reflexes and Somatosensory Conditioning. Journal of Neurophysiology, 91(4), 1516–1523. 10.1152/jn.00695.2003

Gomes, M. M., Jenz, S. T., Beauchamp, J. A., Negro, F., Heckman, C. J., & Pearcey, G. E. P. (2024). Voluntary co-contraction of ankle muscles alters motor unit discharge characteristics and reduces estimates of persistent inward currents. The Journal of Physiology, 602(17), 4237–4250. 10.1113/JP286539

Gorassini, M., Yang, J. F., Siu, M., & Bennett, D. J. (2002a). Intrinsic Activation of Human Motoneurons: Possible Contribution to Motor Unit Excitation. Journal of Neurophysiology, 87(4), 1850–1858. 10.1152/jn.00024.2001

Gorassini, M., Yang, J. F., Siu, M., & Bennett, D. J. (2002b). Intrinsic Activation of Human Motoneurons: Reduction of Motor Unit Recruitment Thresholds by Repeated Contractions. Journal of Neurophysiology, 87(4), 1859–1866. 10.1152/jn.00025.2001

Hassan, A., Thompson, C. K., Negro, F., Cummings, M., Powers, R. K., Heckman, C. J., Dewald, J. P., & McPherson, L. M. (2020). Impact of parameter selection on estimates of motoneuron excitability using paired motor unit analysis. Journal of Neural Engineering, 17(1), 016063.

Heckman, C. J., & Enoka, R. M. (2012). Motor unit. Comprehensive Physiology, 2(4), 2629–2682.

Heckman, C. J., Mottram, C., Quinlan, K., Theiss, R., & Schuster, J. (2009). Motoneuron excitability: The importance of neuromodulatory inputs. Clinical Neurophysiology, 120(12), 2040–2054.

Holobar, A., & Farina, D. (2014). Blind source identification from the multichannel surface electromyogram. Physiological Measurement, 35(7), R143–165. 10.1088/0967-3334/35/7/R143

Holobar, A., Minetto, M. A., & Farina, D. (2014). Accurate identification of motor unit discharge patterns from high-density surface EMG and validation with a novel signal-based performance metric. Journal of Neural Engineering, 11(1), 016008. 10.1088/1741-2560/11/1/016008

Hug, F., Avrillon, S., Del Vecchio, A., Casolo, A., Ibanez, J., Nuccio, S., Rossato, J., Holobar, A., & Farina, D. (2021). Analysis of motor unit spike trains estimated from high-density surface electromyography is highly reliable across operators. Journal of Electromyography and Kinesiology, 58, 102548. 10.1016/j.jelekin.2021.102548

Hyngstrom, A. S., Johnson, M. D., Miller, J. F., & Heckman, C. J. (2007). Intrinsic electrical properties of spinal motoneurons vary with joint angle. Nature Neuroscience, 10(3), Article 3. 10.1038/nn1852

Jacobs, B. L., Martın-Cora, F. J., & Fornal, C. A. (2002). Activity of medullary serotonergic neurons in freely moving animals. Brain Research Reviews, 40(1–3), 45–52.

Jenz, S. T., Beauchamp, J. A., Gomes, M. M., Negro, F., Heckman, C. J., & Pearcey, G. E. P. (2023). Estimates of persistent inward currents in lower limb motoneurons are larger in females than in males. Journal of Neurophysiology, 129(6), 1322–1333. 10.1152/jn.00043.2023

Johnson, M. D., Thompson, C. K., Tysseling, V. M., Powers, R. K., & Heckman, C. J. (2017). The potential for understanding the synaptic organization of human motor commands via the firing patterns of motoneurons. Journal of Neurophysiology, 118(1), 520–531.

Katz, P. S. (2016). Evolution of central pattern generators and rhythmic behaviours. Philosophical Transactions of the Royal Society B: Biological Sciences, 371(1685), 20150057. 10.1098/rstb.2015.0057

Kaupp, C., Pearcey, G. E. P., Klarner, T., Sun, Y., Cullen, H., Barss, T. S., & Zehr, E. P. (2018). Rhythmic arm cycling training improves walking and neurophysiological integrity in chronic stroke: The arms can give legs a helping hand in rehabilitation. Journal of Neurophysiology, 119(3), 1095–1112. 10.1152/jn.00570.2017

Kavanagh, J. J., McFarland, A. J., & Taylor, J. L. (2019). Enhanced availability of serotonin increases activation of unfatigued muscle but exacerbates central fatigue during prolonged sustained contractions. The Journal of Physiology, 597(1), 319–332. 10.1113/JP277148

Khurram, O. U., Negro, F., Heckman, C. J., & Thompson, C. K. (2021). Estimates of persistent inward currents in tibialis anterior motor units during standing ramped contraction tasks in humans. Journal of Neurophysiology, 126(1), 264–274.

Kuo, J. J., Lee, R. H., Johnson, M. D., Heckman, H. M., & Heckman, C. J. (2003). Active Dendritic Integration of Inhibitory Synaptic Inputs In Vivo. Journal of Neurophysiology, 90(6), 3617–3624. 10.1152/jn.00521.2003

Kuznetsova, A., Brockhoff, P. B., & Christensen, R. H. B. (2017). lmerTest Package: Tests in Linear Mixed Effects Models. Journal of Statistical Software, 82, 1–26. 10.18637/jss.v082.i13

Lenth, R. V., Banfai, B., Bolker, B., Buerkner, P., Giné-Vázquez, I., Herve, M., Jung, M., Love, J., Miguez, F., Piaskowski, J., Riebl, H., & Singmann, H. (2025). emmeans: Estimated Marginal Means, aka Least-Squares Means (Version 1.11.1) [Computer software]. https://cran.r-project.org/web/packages/emmeans/index.html

Loadman, P. M., & Zehr, E. P. (2007). Rhythmic arm cycling produces a non-specific signal that suppresses Soleus H-reflex amplitude in stationary legs. Experimental Brain Research, 179(2), 199–208. 10.1007/s00221-006-0782-2

MacDonell, C. W., Power, K. E., Chopek, J. W., Gardiner, K. R., & Gardiner, P. F. (2015). Extensor motoneurone properties are altered immediately before and during fictive locomotion in the adult decerebrate rat. The Journal of Physiology, 593(10), 2327–2342.

Mesquita, R. N. O., Taylor, J. L., Trajano, G. S., Škarabot, J., Holobar, A., Gonçalves, B. A. M., & Blazevich, A. J. (2022). Effects of reciprocal inhibition and whole-body relaxation on persistent inward currents estimated by two different methods. The Journal of Physiology, 600(11), 2765–2787. 10.1113/JP282765

Orssatto, L. B. R., Fernandes, G. L., Blazevich, A. J., & Trajano, G. S. (2022). Facilitation– inhibition control of motor neuronal persistent inward currents in young and older adults. The Journal of Physiology, 600(23), 5101–5117. 10.1113/JP283708

Orssatto, L. B. R., Mackay, K., Shield, A. J., Sakugawa, R. L., Blazevich, A. J., & Trajano, G. S. (2021). Estimates of persistent inward currents increase with the level of voluntary drive in low-threshold motor units of plantar flexor muscles. Journal of Neurophysiology, 125(5), 1746–1754. 10.1152/jn.00697.2020

Pearcey, G. E., Khurram, O. U., Beauchamp, J. A., Negro, F., & Heckman, C. J. (2022). Antagonist tendon vibration dampens estimates of persistent inward currents in motor units of the human lower limb. bioRxiv.

Pearcey, G. E. P., & Zehr, E. P. (2019). Exploiting cervicolumbar connections enhances short-term spinal cord plasticity induced by rhythmic movement. Experimental Brain Research, 237(9), 2319–2329. 10.1007/s00221-019-05598-9

Porter, J. A., Barss, T. S., Mann, D. J., Karamzadeh, Z., Okusanya, D. O., Hemakumara, S. G., Zehr, E. P., Klarner, T., & Mushahwar, V. K. (2025). Pushing the Limits of Interlimb Connectivity: Neuromodulation and Beyond. Biomedicines, 13(5), 1228. 10.3390/biomedicines13051228

Power, K. E., McCrea, D. A., & Fedirchuk, B. (2010). Intraspinally mediated state-dependent enhancement of motoneurone excitability during fictive scratch in the adult decerebrate cat. The Journal of Physiology, 588(15), 2839–2857.

Powers, R. K., Nardelli, P., & Cope, T. C. (2008). Estimation of the Contribution of Intrinsic Currents to Motoneuron Firing Based on Paired Motoneuron Discharge Records in the Decerebrate Cat. Journal of Neurophysiology, 100(1), 292–303. 10.1152/jn.90296.2008

Škarabot, J., Beauchamp, J. A., & Pearcey, G. E. P. (2025). Human motor unit discharge patterns reveal differences in neuromodulatory and inhibitory drive to motoneurons across contraction levels. Journal of Neurophysiology, 134(5), 1429–1444. 10.1152/jn.00249.2025

Škarabot, J., Thomason, H. W., Nazaroff, B. M., Connelly, C. D., Valenčič, T., Ho, M. L., Tyagi, K., Beauchamp, J. A., & Pearcey, G. E. (2025). Training-induced alterations in the modulation of human motoneuron discharge patterns with contraction force. bioRxiv, 2025.06.02.657380. 10.1101/2025.06.02.657380

Stephenson, J. L., Christou, E. A., & Maluf, K. S. (2011). Discharge rate modulation of trapezius motor units differs for voluntary contractions and instructed muscle rest. Experimental Brain Research, 208(2), 203–215. 10.1007/s00221-010-2471-4

Trajano, G. S., Taylor, J. L., Orssatto, L. B. R., McNulty, C. R., & Blazevich, A. J. (2020). Passive muscle stretching reduces estimates of persistent inward current strength in soleus motor units. Journal of Experimental Biology, 223(21), jeb229922. 10.1242/jeb.229922

Udina, E., D’Amico, J., Bergquist, A. J., & Gorassini, M. A. (2010). Amphetamine increases persistent inward currents in human motoneurons estimated from paired motor-unit activity. Journal of Neurophysiology, 103(3), 1295–1303. 10.1152/jn.00734.2009

Veasey, S. C., Fornal, C. A., Metzler, C. W., & Jacobs, B. L. (1995). Response of serotonergic caudal raphe neurons in relation to specific motor activities in freely moving cats. The Journal of Neuroscience: The Official Journal of the Society for Neuroscience, 15(7 Pt 2), 5346–5359. 10.1523/JNEUROSCI.15-07-05346.1995

Wickham, H., Chang, W., Henry, L., Pedersen, T. L., Takahashi, K., Wilke, C., Woo, K., Yutani, H., Dunnington, D., Brand, T. van den, Posit, & PBC. (2025). ggplot2: Create Elegant Data Visualisations Using the Grammar of Graphics (Version 3.5.2) [Computer software]. https://cran.r-project.org/web/packages/ggplot2/index.html

Wilson, J. M., Thompson, C. K., Miller, L. C., & Heckman, C. J. (2015). Intrinsic excitability of human motoneurons in biceps brachii versus triceps brachii. Journal of Neurophysiology, 113(10), 3692–3699. 10.1152/jn.00960.2014

Zehr, E. P., Barss, T. S., Dragert, K., Frigon, A., Vasudevan, E. V., Haridas, C., Hundza, S., Kaupp, C., Klarner, T., Klimstra, M., Komiyama, T., Loadman, P. M., Mezzarane, R. A., Nakajima, T., Pearcey, G. E. P., & Sun, Y. (2016). Neuromechanical interactions between the limbs during human locomotion: An evolutionary perspective with translation to rehabilitation. Experimental Brain Research, 234(11), 3059–3081. 10.1007/s00221-016-4715-4

Zehr, E. P., Carroll, T. J., Chua, R., Collins, D. F., Frigon, A., Haridas, C., Hundza, S. R., & Thompson, A. K. (2004). Possible contributions of CPG activity to the control of rhythmic human arm movement. Canadian Journal of Physiology and Pharmacology, 82(8–9), 556–568. 10.1139/y04-056

